# At least 10 genes on chromosome 5 of *Candida albicans* are downregulated in concert to control cell wall and to confer adaptation to caspofungin

**DOI:** 10.1101/2023.02.06.527048

**Authors:** Sudisht K. Sah, Anshuman Yadav, Elena Rustchenko

**Author notes:** authors contributed equally (Sudisht K. Sah, Anshuman Yadav).

## Abstract

*Candida albicans* is part of normal microbiota, however, can cause superficial and life threatening infection in immune-compromised individuals. Drugs from echinocandin (ECN) class that disrupt cell wall synthesis, are being used as a major treatment strategy against candidiasis. As the use of ECNs for the treatment of candidiasis is increasing, resistance against ECNs is also emerging. Previously, we reported involvement of 5 chromosome 2 (Ch2) genes in adaptation to ECN drugs. Here, we explored 22 candidate-genes on Ch5 that are consistently downregulated in independent mutants adapted to caspofungin (CAS), for their role in ECN adaptation. We also compared cell wall remodelling in CAS-adapted mutants and in 10 knockouts (KOs) from Ch5. Independent KO experiments as combined with broth microdilution assay, demonstrated that, as expected, 10 out of 22 Ch5 genes decrease ECN susceptibility by controlling the levels of three major components of the cell wall, glucan, mannan, and chitin. Some KOs decreased glucan or increased chitin or both. Similar cell wall remodelling, decreased glucan and increased chitin, was found in CAS-adapted mutants with no ploidy change. Some other KOs had no glucan change, but increased the level of either mannan or chitin. Our results identify the function of two uncharacterized genes, orf19.970 and orf19.4149.1, and expand the functions of *DUS4, RPS25B, UAP1, URA7, RPO26, HAS1*, and *CKS1*. The function of *CHT2*, as negative regulator of ECN susceptibility, has been previously established. Importantly, half of the above genes are essential indicating that essential processes are involved in cell wall remodelling for adaptation to ECNs. Also important, orf19.970 and orf19.4149.1 have no human orthologues. Finally, our work shows that multiple mechanisms are used by *C. albicans* cells to remodel cell wall in order to adapt to CAS. This work continues to identify common pathways that are involved in drug adaptation, as well as new genes controlling ECN susceptibility and reveals new targets for development of novel antifungal drugs.

## INTRODUCTION

*Candida albicans* is an opportunistic unicellular fungus that lives as normal microbiota in human gut and genital organs (1, 2). Candidiasis caused by *C. albicans* along with other non-albicans species is emerging as a major problem worldwide (3). As echinocandins (ECNs) are being used as frontline treatment against candidiasis, there is slow, but consistent increase in ECNs resistance in *Candida* species (1). The only known mechanism of clinical resistance to ECNs in *C. albicans* is point mutations in *FKS1* (orf19.2929) which encodes for the catalytic subunit of 1,3-*β*-glucan synthase complex (4). However, there are mechanisms which can decrease or increase ECNs susceptibility without any mutations in *FKS1* gene (reviewed in 5). For example, several laboratories published genes of *C. albicans* and the related species *C. glabrata* that can increase or decrease ECNs susceptibility (reviewed in 6). Here, we will use the term “tolerance” or “adaptation” referring to a cellular condition that allows cells to survive at or below the MIC (Minimum Inhibitory Concentration) breakpoint without forming mechanism-specific *FKS1* resistance mutations (7).

We have previously generated caspofungin (CAS)-adapted mutants by a direct exposure of *C. albicans* cells to CAS on Petri dishes in independent parallel experiments (8). The adapted mutants had 2-8 fold increased MICs in the absence of *FKS1* classic mutations to resistance. Adaptation depended on different mechanisms including various aneuploidies of chromosome 5 (Ch5), as well as a still unknown mechanism, which is independent of aneuploidy (8). We initiated the search for the genes that decrease ECN susceptibility for adaptation, for which we performed RNA-seq with two no ploidy change mutants and three mutants that acquired Ch5 monosomy. We compared differentially expressed gene (DEG) in mutants vs parentals and across all comparisons. Importantly, non-random distribution of DEG was identified: most of upregulated genes reside on Ch2, whereas most of downregulated genes reside on Ch5 (6). The non-random distribution is consistent with the previously demonstrated role of ploidy of these chromosomes in ECN susceptibility, i.e. trisomy of Ch2 or monosomy of Ch5 conferring tolerance to CAS (5). 5 positive regulators on Ch2 were previously validated and shown to control three major components of the cell wall, glucan, mannan, and chitin (6). In this work, we have validated and characterized a set of 10 Ch5 genes that serve as negative regulators of ECN susceptibility and also control the amount of glucan, mannan and chitin in cell wall.

## RESULTS AND DISCUSSION

### Multiple Ch5 genes serve as negative regulators of drug susceptibility in adaptation to CAS

As presented in the Introduction, our search of genes that have a role in ECN susceptibility, identified 26 genes downregulated across five CAS-adapted mutants (5). Of 26 genes, 22 were dispersed across Ch5 (Fig 1A). We used transient CRISPR/Cas9 method to independently delete each of 22 genes on Ch5 in CAF4-2 background (Table 1) (see also Materials and Methods). The knockouts (KOs) were tested for their role in ECN susceptibility with broth microdilution assay (Materials and Methods) in the presence of CAS. 12 out of 22 KOs could not be validated with this approach. However, 10 KOs lacking *CHT2, DUS4, RPS25B, UAP1*, orf19.970, or a single copy of essential *URA7, RPO26, HAS1, CKS1*, and orf19.4149.1, showed, as expected, increased growth vs parental strain (Fig. 1B). These KOs acquired 2-4-fold increase of CAS MIC90. On a special note, confirming *CHT2* role as a negative regulator of ECN susceptibility (9) validates our approach.

**FIG 1.**
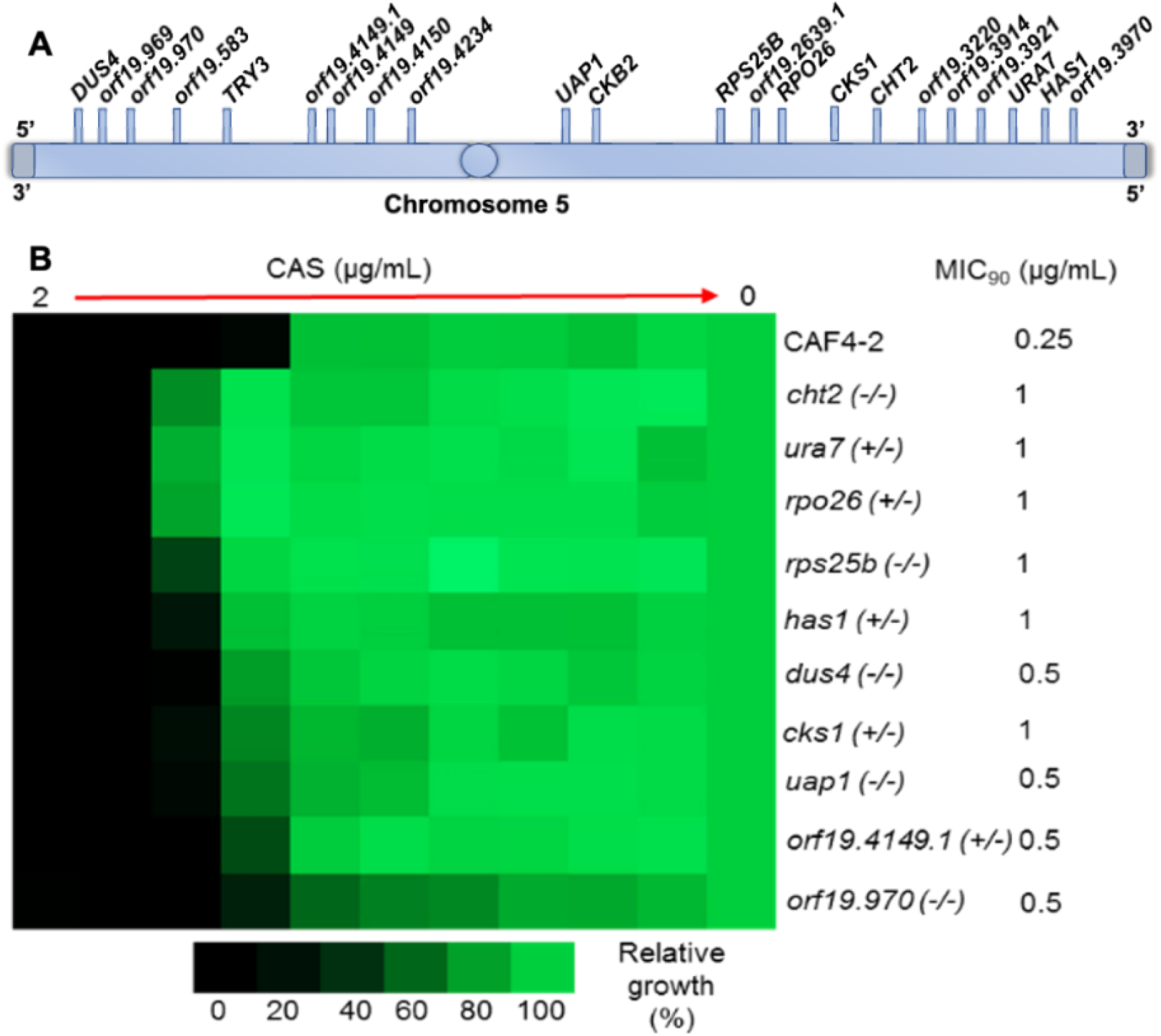
Validation of Ch5 KOs with broth microdilution assay. **A.** A cartoon presenting the distribution of 22 downregulated genes across Ch5 (not to scale). **B.** The heat map shows the increased growth of Ch5 KOs vs their parental CAF4-2. CAS refers to caspofungin. Shown are the parental strain CAF4-2 and 10 independent mutants lacking both copies of *CHT2, DUS4, RPS25B, UAP1* and orf19.970, as well as one copy of essential *URA7, RPO26, HAS1, CKS1*, and orf19.4149.1. Names, genotypes of strains and MIC90 are indicated on the right. The assay was conducted according to CLSI methods in RPMI 1640 medium with 2% glucose. The assay included a maximum caspofungin concentration of 2 μg/mL and 2-fold serial dilutions. A total of 10^3^ cells were inoculated into each well in four technical replicates and the tray was incubated at 35°C for 24 h. Control wells without the drug or without cells were included. The no-cell control was used to subtract the background. The no-drug control was used for normalization. The colour bar for percent growth is presented below the heap map.

**Table 1.** *C. albicans* strains and CAS-adapted mutants used in the study. Adapted and modified from Sah et al., 2022 (4).

| Strains | Description | Phenotype | Source and/or reference |
| --- | --- | --- | --- |
| JRCT1 | Parental strain; diploid | Adapted to caspofungin | (8) |
| JMC200-2-5 | Same as JRCT1, but see the phenotype | Same as above | ( 8) |
| JMC160-2-5 | Same as JRCT1, but see the phenotype | Same as above | ( 8) |
| SC5314 | Parental strain; diploid | Susceptible to caspofungin | (20) |
| SMC60-3-4 | Same as SC5314 but with Ch5 monosomy; *MTLa* | Adapted to caspofungin | ( 8) |
| SMC60-2-5 | Same as SC5314 but with Ch5 monosomy; *MTLa* | Same as above | ( 8) |
| CAF4-2 | Ura<sup>-</sup> derivative of SC5314 | Susceptible to caspofungin | ( 21) |

### Susceptibility of KOs from Ch5 to calcofluor white and zymolyase

By testing 10 Ch5 validated KOs against calcofluor white, we found no cell wall disturbance in 6 KOs (Fig. 2A). Consistently, these KOs and, intriguingly, *rps25b* that showed cell wall disturbance in the presence of calcofluor white, were lysed by zymolyase significantly slower, as compared to the parental strain CAF4-2 (Fig. 2B). As essential activities of zymolyase include β-1,3-glucan laminaripentao-hydrolase activity and β-1,3-glucanase activity (10), our data imply decreased surface exposure of cell wall glucan in the above 7 KOs. Of 3 remaining KOs that did not grow when exposed to calcofluor white (Fig. 2B), two, *dus4* and *rpo26*, were, consistently, also sensitive to zymolyase (Fig. 2B), whereas orf19.970 had no change in zymolyase sensitivity, as compared to the parental strain. Our data indicate multiple mechanisms by which Ch5 negative regulators govern cell wall.

**FIG 2.**
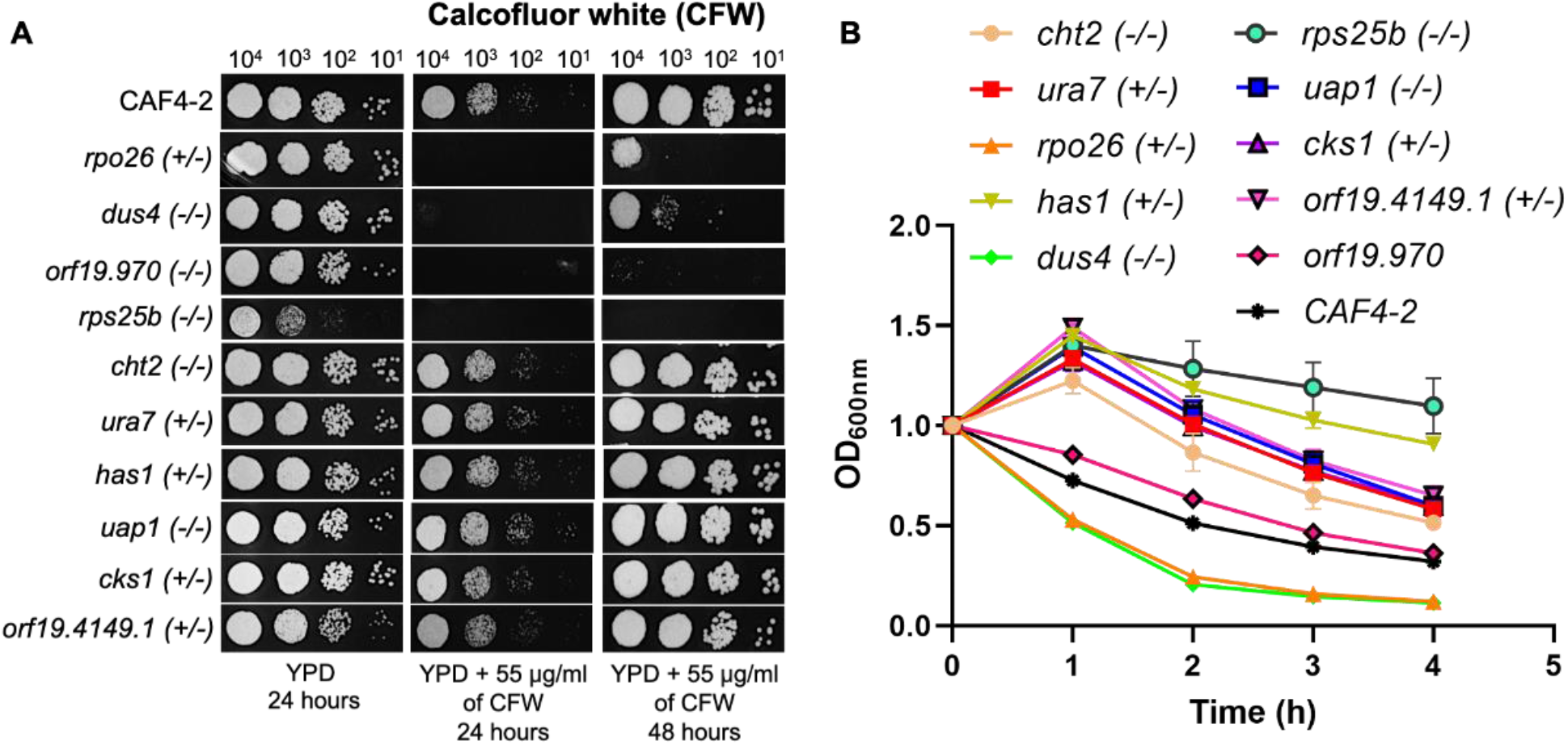
Survival of 10 KOs from Ch5 vs parental strain CAF4-2 in the presence of calcofluor white (CFW) or zymolyase. **A.** Spot assay shows growth of 10 KOs lacking both copies of *CHT2, DUS4, RPS25B, UAP1* and orf19.970, or one copy of essential *URA7, RPO26, HAS1, CKS1* and *orf19.4149.1* vs the parental strain CAF4-2 on YPD medium supplemented with 55 μg/mL of CFW. Strains, genotypes, cell number spotted and incubation time of plates, as well as control growth on YPD medium are shown. **B.** Survival in the presence of zymolyase of 10 KOs and parental strain as mentioned in **A**. The means and standard deviations of the optical density (O.D_600nm_) values of each strain from four technical replicates are plotted against the time of incubation with zymolyase. The O.D_600nm_ at time point zero is scaled to 1 for better representation. Student’s *t* test was performed between the parental strain and each mutant strain at the 4-h time points. *P* values for the time point ranged from 0.001055 to 1.77*10^−7^, except for orf19.970 where the *P* value was 0.076.

### Remodelling of cell wall is key to CAS adaptation

We set about to compare amounts of three major components of cell wall, glucan, mannan, and chitin, in four CAS-adapted mutants (Figs. 3A and B) and 10 KOs from Ch5 (Figs. 4A and B). Interestingly, CAS-adapted mutants with no ploidy change showed downregulation of glucan and mannan, two components of pathogen associated molecular pattern (PAMP). These were accompanied by twice increase of the amount of chitin, which is believed to fortify the cell wall (11). It is important that decrease of glucan also occurred in 7 out of 10 KOs, while remaining three KOs had no glucan change. Furthermore, increase of chitin occurred in 5 out of 10 KOs, while remaining five KOs had no chitin change (Figs. 4A and B). It seems that despite apparent complexities involved in cell wall control, there are some underlying similarities between cell wall condition of CAS-adapted mutants with no ploidy change and some Ch5 negative regulators. Decrease of glucan and increase of chitin are also produced, when Ch2 genes are upregulated, as can be deduced from the patterns of their KOs (6). Note that KO of positive regulator produces opposite result to the genes’ action in cell. Thus, upregulation of Ch2 genes can also be responsible for decrease of glucan and increase of chitin in cells with no ploidy change. However, cell wall remodelling in CAS-adapted mutants with Ch5 monosomy that features increase of all three components, glucan, mannan, and chitin (Fig. 3), is different from adapted mutants with no ploidy change and can be explained only partially by the action of genes on Ch2 and Ch5 that confer increase of chitin. Presumably, the absence of one copy of Ch5 activates some additional factors that contribute to adaptation, and act to partially override the action of Ch2 and Ch5 genes.

**FIG 3.**
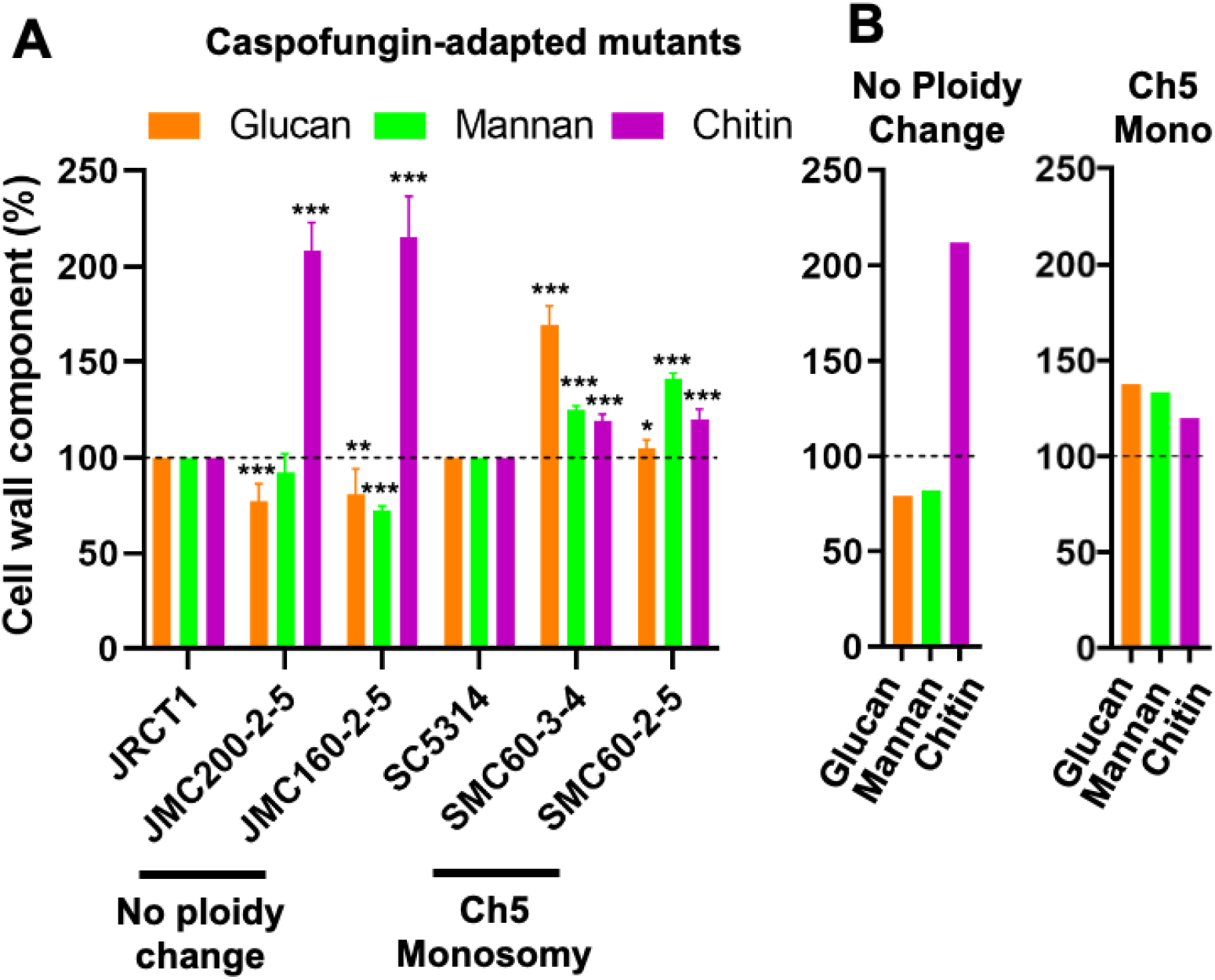
CAS-adapted mutants exhibit altered levels of cell wall mannan, glucan, and chitin, as compared to the parental strains. **A**. Shown are parental strains JRCT1 and SC5314, mutants with no ploidy change or with Ch5 monosomy, as indicated. Measurements were performed on two biological replicates with four technical replicates for glucan and chitin and one biological replicate with five technical replicates for mannan. The amount of mannan, glucan and chitin in the parental strains is set as 100%. The asterisks indicate a p-value of <0.05 (*) or <0.01 (**) or <0.001 (***), determined using Student’s *t* test. **B**. Cartoons showing average amounts of glucan, mannan and chitin for the CAS-adapted mutants presented in **A**. Mutants with no ploidy change and with Ch5 monosomy are indicated.

**FIG 4.**
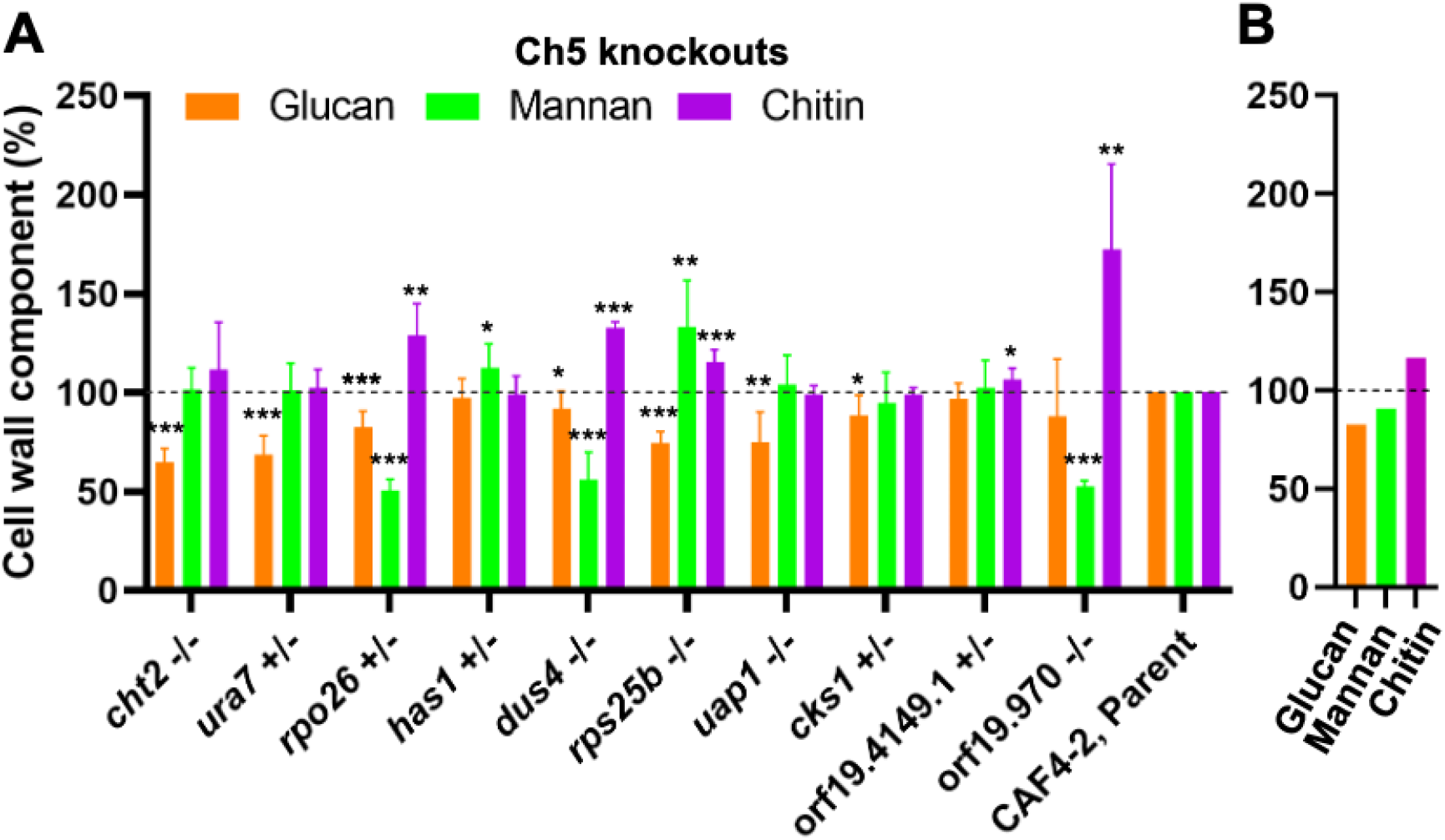
KOs from Ch5 exhibit altered levels of cell wall mannan, glucan and chitin. **A**. The panel shows parental strain CAF4-2 and 5 independent KOs lacking both copies of *CHT2, DUS4, RPS25B, UAP1* or orf19.970, and 5 KOs lacking one copy of *URA7, RPO26, HAS1, CKS1* or orf19.4149.1. Measurements were performed on two biological replicates with four technical replicates for glucans, mannans and chitin. The amount of mannan, glucan and chitin in the parental strain is set as 100%. The asterisks indicate a *P* value of <0.05 (*) or <0.01 (**) or <0.001 (***), determined using Student’s *t* test. **B.** A cartoon presenting average amounts of glucan, mannan and chitin for the strains presented in A.

### Properties of proteins encoded by the 10 simultaneously downregulated Ch5 genes

We performed the Gene Ontology (GO) and Kyoto Encyclopedia of Genes and Genomes (KEGG) and analysed the 35 similarly expressed genes in CAS-adapted mutants that included 10 genes from Ch5 of our current study and we found no significant enrichment of any biological pathway (6).

In order to understand molecular function of proteins that are encoded by ten simultaneously downregulated genes on Ch5, we explored Candida Genome Database for the available information. The size of proteins ranges from 102 aa to 583 aa (amino acid). We could not find any common motifs or domains present when we did multiple sequence alignment of those genes.

*CHT2* is well-characterized gene encoding for GPI-linked chitinase. *URA7* is 44% identical to *Saccharolobus solfataricus* cytidine triphosphate (CTP) synthase. *RPO26* encodes for a subunit of RNA polymerase and is 67 % identical to *Saccharomyces cerevisiae* Rpo26p. *HAS1* is functional homolog of *S. cerevisiae HAS1*, which encodes for nucleolar protein of the DEAD-box-ATP dependent RNA helicase and is involved in ribosome biogenesis. *DUS4* is involved in tRNA-dihydrouridine synthesis and shares 29% identity with human homolog of tRNA-dihydrouridine synthase catalytic domain. *RPS25B* is a ribosomal protein sharing 85% identity with *S. cerevisiae* Rps25bp, which serves as structure component of ribosome. *UAP1* encodes for UDP-N-acetylglucosamine pyrophosphorylase, which catalyses synthesis of UDP-N-acetylglucosamine. *CKS1* shares 86 % identity with *S. cerevisiae* cell cycle regulatory protein Cks1p. The molecular function of Cks1p is cyclin-dependent protein serine/threonine kinase activator activity. Orf19.4149.1 encodes for a protein component of small (40S) ribosomal subunit which is identical to 40S-eif1a from *Kluyveromyces lactis*. The putative function of orf19.970 is poorly understood, except that it has a role in microtubule related process and shares 30 % sequence identity with *S. cerevisiae BER1*.

### Conclusions

We find that cell wall of CAS-adapted mutants with no ploidy change (decrease of glucan and mannan and increase of chitin) is remodelled differently than that of CAS-adapted mutants with Ch5 monosomy (increase of all three components). In this regard, it worth to mention that adapted mutants with no ploidy change differ from mutants with Ch5 monosomy not only by their cell wall remodelling, but also by their pattern of adaptation to three ECNs, CAS, ANI, and MFG. The former are adapted to all three ECNs, whereas the latter are adapted to CAS only (12). We continue identifying common pathways in different CAS-adapted mutants by finding that 10 Ch5 genes that are downregulated in both classes of adapted mutants have a role in ECN susceptibility as negative regulators, and act in concert to remodel the cell wall for adaptation to CAS. Half of these genes seem to increase the level of chitin consistent with the cell wall salvage concept (11) and similar to chitin increase, as observed in both classes of CAS-adapted mutants. Also, all but three genes seem to act to decrease the level of glucan, a major component of cell wall and an important component of PAMP, which is consistent with glucan decrease in CAS-adapted mutants with no ploidy change. Interestingly, three remaining genes seem to represent an alternative mechanism(s) of adaptation to ECNs by which adaptation occurs with no change of glucan level. Intriguingly, while many Ch5 genes seem to contribute directly to the pattern of cell wall remodelling in CAS-adapted mutants with no ploidy change, the contribution of these genes in the pattern of cell wall remodelling of adapted mutants with Ch5 monosomy is not as obvious. Further studies are needed to detail how two cohorts of genes, positive regulators from Ch2 (6) and negative regulators from Ch5 control adaptation to ECNs by cell wall remodelling.

## MATERIALS & METHODS

### Strains, plasmids, and primers

We used four *C. albicans* CAS-adapted mutants that arose due to independent mutational events from two genetic backgrounds *i.e*. SC5314 and the clinical isolate JRCT1 (Table 1). The chromosome conditions of all strains were extensively characterized previously (8) and confirmed by DNA sequencing (DNA-seq), which will be presented elsewhere.

The primers and plasmids used in this study are presented in Table S1 in the supplemental material. Maintenance and growth of strains and media. Cells were maintained, stored, and grown using our standardized approach that prevents the induction of chromosome instability, as previously described (13). This approach favors maintaining cells that represent a major fraction of the population of cells (14). Briefly, cells were stored at −80°C. When needed, cells from a −80°C stock were streaked for independent colonies onto yeast extract-peptone-dextrose (YPD) plates and incubated at 37°C until young colonies with a size of approximately 1*10^5^ to 3*10^5^ cells/colony grew up. Young colonies were collected, a proper dilution in sterile water was prepared with the aid of a hemacytometer, approximately 3,000 CFU were plated onto each plate, and plates were incubated until young colonies appeared.

Cells were stored in a 25% (vol/vol) glycerol solution at −80°C to interrupt metabolism and routinely grown at 37°C.

YPD medium (1% yeast extract, 2% peptone, 2% dextrose). RPMI 1640 medium (Sigma, St. Louis, MO, USA) was supplemented with 2% glucose. In order to prepare solid medium, 2% (wt/vol) agar was added. Nourseothricin at 150 mg/mL (Jena Bioscience GmbH, Jena, Germany), uridine at 50 mg/mL (Sigma-Aldrich, St. Louis, MO, USA), CAS (Merck Sharp & Dohme Corp., Kenilworth, NJ, USA), or anidulafungin (Pfizer Inc., New York, NY, USA) was added when needed. We also used Zymolyase 100T (U.S. Biological, Swampscott, MA, USA).

### Gene deletions by the transient CRISPR method

We used the transient CRISPR-CAS9 method allowing the deletion of both copies of the target genes in *C. albicans* with a single transformation (15). The *C. albicans* cells were transformed by lithium acetate transformation method (9). NAT^R^ transformants were selected on YPD plates supplemented with 150 mg/mL of nourseothricin. The correct target gene deletion was verified by PCR amplification using the 59-flanking region and either the gene of interest (GI), NAT^R^ as well as by the amplification of the entire GI or NAT^R^, using 59- and 39-flanking regions (6).

### Broth microdilution assay to determine MICs

We employed a broth microdilution assay according to the CLSI document M27-A3 broth microdilution method for yeasts (16), with some modifications (6). The turbidities were measured with a Spark multimode micro-plate reader (Tecan, Zurich, Switzerland) at 600 nm. Normalized readings were generated in Microsoft Excel and presented as heat maps.

### Zymolyase assay

Cells were streaked from a −80°C stock onto YPD plates for independent colonies and incubated at 37°C for 19 h. Colonies were then collected, and cells were counted with a hemacytometer. The assay was conducted as described in paper (6).

### Determination of glucan content in the cell wall

The level of bulk of glucans was determined by an aniline blue assay (17, 6).

### Determination of chitin content in the cell wall

The amount of chitin was determined by measuring the absorbance of glucosamine released by acid hydrolysis of the purified cell wall as described previously (18, 6).

### Determination of mannan content in the cell wall

Mannan contents were determined with the alcian blue staining method (19, 6).

## ACKNOWLEDGEMENT

We thank Merck and Co., Inc., for the generous donation of caspofungin and Pfizer Inc., for the generous donation of anidulafungin via a compound transfer program.

## AUTHOR CONTRIBUTIONS

Conceptualization: SKS, AY, ER

Methodology: SKS, AY

Investigation: SKS, AY, ER

Visualization: SKS, AY

Funding acquisition: ER

Supervision: ER

Writing – original draft: SKS, AY, ER

Writing – review & editing: SKS, AY, ER

## DATA AVAILABILITY

All data are available in the main text or supplemental materials.

## SUPPLEMENTAL MATERIALS

**Table S1.** List of primers and plasmids used in the study

**FIG S1.** Growth curves of *C. albicans* deletion mutants vs parental CAF4-2.

