## Supplemental Material for "At least 10 genes on chromosome 5 of *Candida albicans* are downregulated in concert to control cell wall and to confer adaptation to caspofungin"

**SUPPLEMENTAL MATERIALS**

Table S1. List of primers and plasmids used in the study

| **Primer purpose** | **Gene** | **Primer name and sequence** |
| --- | --- | --- |
| Amplification of *CaCas9* cassette | *CaCas9* | CaCas9 F, ATCTCATTAGATTTGGAACTTGTGGGGTT CaCas9 R, TTCGAGCGTCCCAAAACCTTCT |
| Amplification of SNR52 promoter and sgRNA scaffold |  | SNR52 F, AAGAAAGAAAGAAAACCAGGAGTGAA sgRNA R, ACAAATATTTAAACTCGGGACCTGG |
| Amplification of final sgRNA expression cassette |  | SNR52/N F, GCGGCCGCAAGTGATTAGACT sgRNA/N R, GCAGCTCAGTGATTAAGAGTAAAGAT GG |
| Amplification of sgRNA promoter and *SNR52* scaffold with overlapping ORF/gene guide sequence | *CHT2*  (orf19.3895) | sgRNA/F_19.3895, GGTGTTAGAACCTGAACCAGGTTTTAGAGCTAGAAATAGCAAGTTAAA  SNR52/R_19.3895, CTGGTTCAGGTTCTAACACCCAAATTAAAAATAGTTTACGCAAGTC |
| Same as above | *URA7*  (orf19.3941) | SNR52/R_19.3941, TCGGTGACATTGAAAGTGCTCAAATTAAAAATAGTTTACGCAAGTC  sgRNA/F_19.3941, AGCACTTTCAATGTCACCGAGTTTTAGAGCTAGAAATAGCAAGTTAAA |
| Same as above | *RPO26*  (orf19.2643) | SNR52/R_19.2643, CCATTTGCTGATGGATCGTCCAAATTAAAAATAGTTTACGCAAGTC  sgRNA/F_19.2643, GACGATCCATCAGCAAATGGGTTTTAGAGCTAGAAATAGCAAGTTAAA |
| Same as above | *RPS25B*  (orf19.6663) | SNR52/R_19.6663, CCGCTTTAGCTGGTGGTAAACAAATTAAAAATAGTTTACGCAAGTC  sgRNA/F_19. 6663, TTTACCACCAGCTAAAGCGGGTTTTAGAGCTAGAAATAGCAAGTTAAA |
| Same as above | *HAS1*  (orf19.3962) | SNR52/R_19.3962, CAGGGAGATTATTGGATCATCAAATTAAAAATAGTTTACGCAAGTC  sgRNA/F_19. 3962, ATGATCCAATAATCTCCCTGGTTTTAGAGCTAGAAATAGCAAGTTAAA |
| Same as above | *DUS4*  (orf19.966) | SNR52/R_19.966, CAGTTGATACCTATACCATCCAAATTAAAAATAGTTTACGCAAGTC  sgRNA/F_19. 966, GATGGTATAGGTATCAACTGGTTTTAGAGCTAGAAATAGCAAGTTAAA |
| Same as above | *CKS1*  (orf19.1282) | SNR52/R_19.1282, CACGATTACTTTAATCAAGACAAATTAAAAATAGTTTACGCAAGTC  sgRNA/F_19.1282, TCTTGATTAAAGTAATCGTGGTTTTAGAGCTAGAAATAGCAAGTTAAA |
| Same as above | *UAP1*  (orf19.4265) | SNR52/R_19.4265, TTGGGAGCACTTGAACCTAACAAATTAAAAATAGTTTACGCAAGTC  sgRNA/F_19.4265, TTAGGTTCAAGTGCTCCCAAGTTTTAGAGCTAGAAATAGCAAGTTAAA |
| Same as above | orf19.4149.1 | SNR52/R_19.4149.1, TTCGTCATAGTATTTCATTCCAAATTAAAAATAGTTTACGCAAGTC  sgRNA/F_19.4149.1, GAATGAAATACTATGACGAAGTTTTAGAGCTAGAAATAGCAAGTTAAA |
| Same as above | orf19.970 | SNR52/R_19.970, CCATGCATCCTTGGAACTTACAAATTAAAAATAGTTTACGCAAGTC  sgRNA/F_19.970, TAAGTTCCAAGGATGCATGGGTTTTAGAGCTAGAAATAGCAAGTTAAA |
| Amplification of ORF/gene deletion cassette with NAT1 | *CHT2*  (orf19.3895) | Orf19.3895F,  TCTTCAAATCCAATTTAGTCTTTCGTTCTTATTTAATTTTATACCCTTAACAAAACAAGCCAAAAAGAAATAAAAGTATCGATGATTCATCCCATTCATTCCATC  Orf19.3895R, GTCTTAATAACTATTTGAGGGTTTTTTTTATATATCAATACAAAAAGAAAAAATATTAGAGTAAACAAGAGGTTAATTCAGCCGCTCTAGAACTAGTGGATCT |
| Same as above | *URA7*  (orf19.3941) | Orf19.3941F, TATTTTTTTGTATCAACGAATCAACAATAACAACAATAAGTACTTTTTAATTCCCTCCATCTCTCTATTAGCAAGTAATCGATGATTCATCCCATTCATTCCATC  Orf19.3941R, CATTACTACCTTATATACACATGACAATTAAGGAATAAAATCAATAAAAAAAAACAGCCAAAAGGAAATCACTTACTAATGCCGCTCTAGAACTAGTGGATCT |
| Same as above | *RPO26*  (orf19.2643) | Orf19.2643F, ATCTATCATCAACAAAAACCTCAAATCATCTAACTAATTCCGCCTCCCCCCCTTTATAATTTCAATACAAAACTTACATAGATGATTCATCCCATTCATTCCATC  Orf19.2643R, TCTCTTGGGTTAACTTGGTCTCCTTTTTCTTATTCCTTTAACAATTCGTCTACTCCTGGGTTAATCAATAGTTCTATCGTGCCGCTCTAGAACTAGTGGATCT |
| Same as above | *RPS25B*  (orf19.6663) | Orf19. 6663F, GAATCAGTTATTCCCCTACGGCAGTGAATGAAAAATTCTTATTCAAGTCTAAGTATACTAACTAGTACAATTTTTTTAGAGATGATTCATCCCATTCATTCCATC  Orf19. 6663R, TCATTTATTCTTTAGTAACATTAATAAAGAACAAGATAAATTATAAAATAACCATTAGATATAGATATTTTTTAATTAATGCCGCTCTAGAACTAGTGGATCT |
| Same as above | *HAS1*  (orf19.3962) | Orf19. 3962F, ATTTTTTTTTCCAGGCTTTAGTTGATGAGATGACCTGATACATCTTACTCTTCACAACTGTTTCTTACCAACAACAAAAAGATGATTCATCCCATTCATTCCATC  Orf19. 3962R, GGAATATGGTACTGTTGATACAATGAATTATCACTATTATATTTCTATATAAAGATATGACCTATTAAAACTTATGAGTCGCCGCTCTAGAACTAGTGGATCT |
| Same as above | *DUS4*  (orf19.966) | Orf19. 966F, TAAGATGAGCTTAAATATGGGAGAAAAAAGAAATCTACATTTTTTTTTTCATCACATCAGGTTAATACCAACTACCAAAAGATGATTCATCCCATTCATTCCATC  Orf19. 966R, TTAACAACAAGACTAGTAGTGTTTCACTAGCAAATATAGAAATGCATTAAATTAGTTATAGAAATTAAAAACAAAAGTACGCCGCTCTAGAACTAGTGGATCT |
| Same as above | *CKS1*  (orf19.1282) | Orf19. 1282F, CAATTAGTTCTTTTTTTTCATTTGTTTCCAGAGTTTAGGAAGACTACCATTTTACAATTTTCAATTCAAATATTTTCCCAGATGATTCATCCCATTCATTCCATC  Orf19. 1282R, CTATTTCTTTCCTTTTTATGTTTTAATTCCAGTGTTAACTTTAAAACCGATTCTATAGTATTCAAATATAGTTAATCTTTGCCGCTCTAGAACTAGTGGATCT |
| Same as above | *UAP1*  (orf19.4265) | Orf19. 4265F, CCAACAAAACTAGTTTTATTAATATTGCTGAAATAGAAAGTAAAAGAAATCAGCAACTTCTTACTATTATTATTATTATCGATGATTCATCCCATTCATTCCATC  Orf19. 4265R, TATATAATAAATAGCATAGCGCACATTATATTATATTATATTATATTATATTATGACTTTAGTAACTAAAAAAGTTTACAGCCGCTCTAGAACTAGTGGATCT |
| Same as above | orf19.4149.1 | Orf19.4149.1F, AACCAAAAAAAAAAAATAACTGAAATTTTTTTCAATAGTATTAGAGTATATTCTTAGACTGTATCGAGCATTAAGACACCGATGATTCATCCCATTCATTCCATC  Orf19.4149.1R, AAGAACCTTCAATGGCAGCTTTTGGGGTTTTGAAACCTAAACCAACATCTTTATACCATCTCTTGGTTTTGCCGCTCTAGAACTAGTGGATCTCTTGTTGGCT |
| Same as above | orf19.970 | Orf19.970F, AATCTAACTCCTCCAATAATAGTTATACATAATATAATTTTACAAAACCTTCTTTTTTTTTTTAAAGTTTATTAACAACTGATGATTCATCCCATTCATTCCATC  Orf19.970R, GCAATATTGTCGAAGCATTTATTAGATCCTAAAGAAAATAAGCAAGATAATGAACCTACGGTAAAGAAACAGAAATTGAGGCCGCTCTAGAACTAGTGGATCT |
| Confirmation of deletion of ORF/gene | *CHT2*  (orf19.3895) | Orf19.3895-fwd, TTGATCCAAGATATCTCCAAACTTG  Orf19.3895-rev, GGTAACAGCAGAGGAAGTAGT  Flk19.3895-rev, TGAAGAACAATAACAAACCCATGTA |
| Same as above | *URA7*  (orf19.3941) | Orf19.3941-fwd, ATCCTACTTTAATAATACCGTTTGTTTATTCAC  Orf19.3941-rev, ATGGATTAATGCAAAATTTTCGTTACC  Flk19.3941-rev, GAGGAATGTAAAATTGAAATTACTCTATAATCC |
| Same as above | *RPO26*  (orf19.2643) | Orf19.2643-fwd, TGATTGCAAACTGGTAGTCG  Orf19.2643-rev, CTAGTTCCCAATATTCTTGCTCTTTC  Flk19.2643-rev, CATTGATTAAATACAATACAAAAGGGG |
| Same as above | *RPS25B*  (orf19.6663) | Orf19. 6663-fwd, CAACAACAGTAATAGAAGTGACCAG  Orf19. 6663-rev, TCTGGCTAAAGAACCACCAAT  Flk19. 6663-rev, AAGGGAGATGTTGAATCAGGAA |
| Same as above | *HAS1*  (orf19.3962) | Orf19. 3962-fwd, TCACCTCCCATATTACAGATGTTG  Orf19. 3962-rev, CAATAATTCAATTGCAGGAATTAAGAATGC  Flk19. 3962-rev, AGCAAAGATGATTTACGTGAACG |
| Same as above | *DUS4*  (orf19.966) | Orf19. 966-fwd, GAGAAATTAACATAAGTGGGAGCTAC  Orf19. 966-rev, AACACCGTCTACACCTGTGTA  Flk19. 966-rev, ATTGTGGCCAATTACATCAAGG |
| Same as above | *CKS1*  (orf19.1282) | Orf19. 1282-fwd, GATAGATATCAATTACTAATTTACCCTTGTTTTTTAC  Orf19. 1282-rev, CTGGAGCATGAGTTTCGTAATG  Flk19. 1282-rev, CAATGAGATTGGCTTATAAATAGTGTATAC |
| Same as above | *UAP1*  (orf19.4265) | Orf19.4265-fwd, CTTCCAACTAAACAACAACAAACAAC  Orf19.4265-rev, TTTGATGCCCTTAGAATTCAAATCATC  Flk19.4265-rev, GTTACTGTTGAAAAGATATATGAAAAATTTAGAG |
| Same as above | orf19.4149.1 | Orf19.4149.1-fwd, GGTACACACAAACCAAGTGA  Orf19.4149.1-rev, TTTAGGATGAAGAATATCTTTTTAAACTCC  Flk19.4149.1-rev, TCTACATTGACCAACGGTGA |
| Same as above | orf19.970 | Orf19.970-fwd, ATTGAGAGAGAAAATCGACTACAA  Orf19.970-rev, ATTTAGCACTTCCTCAGTAAGTTC  Flk19.970-rev, ATTTGCAGAATGGTATGTCGGA |
| Confirmation of deletion of ORF/gene | NAT1 | NAT1 RP4, CACAGACGCGTTGAATTGT |
| **Plasmids** | **Description and Purpose** | |
| pV1093 | CaCas9/gRNA cassette carrying ampicillin resistance gene,  amplification of CaCas9/gRNA cassette | |
| pJK863 | CaNAT1-FLP cassette carrying nourseothricin resistance gene, amplification of NAT cassette | |

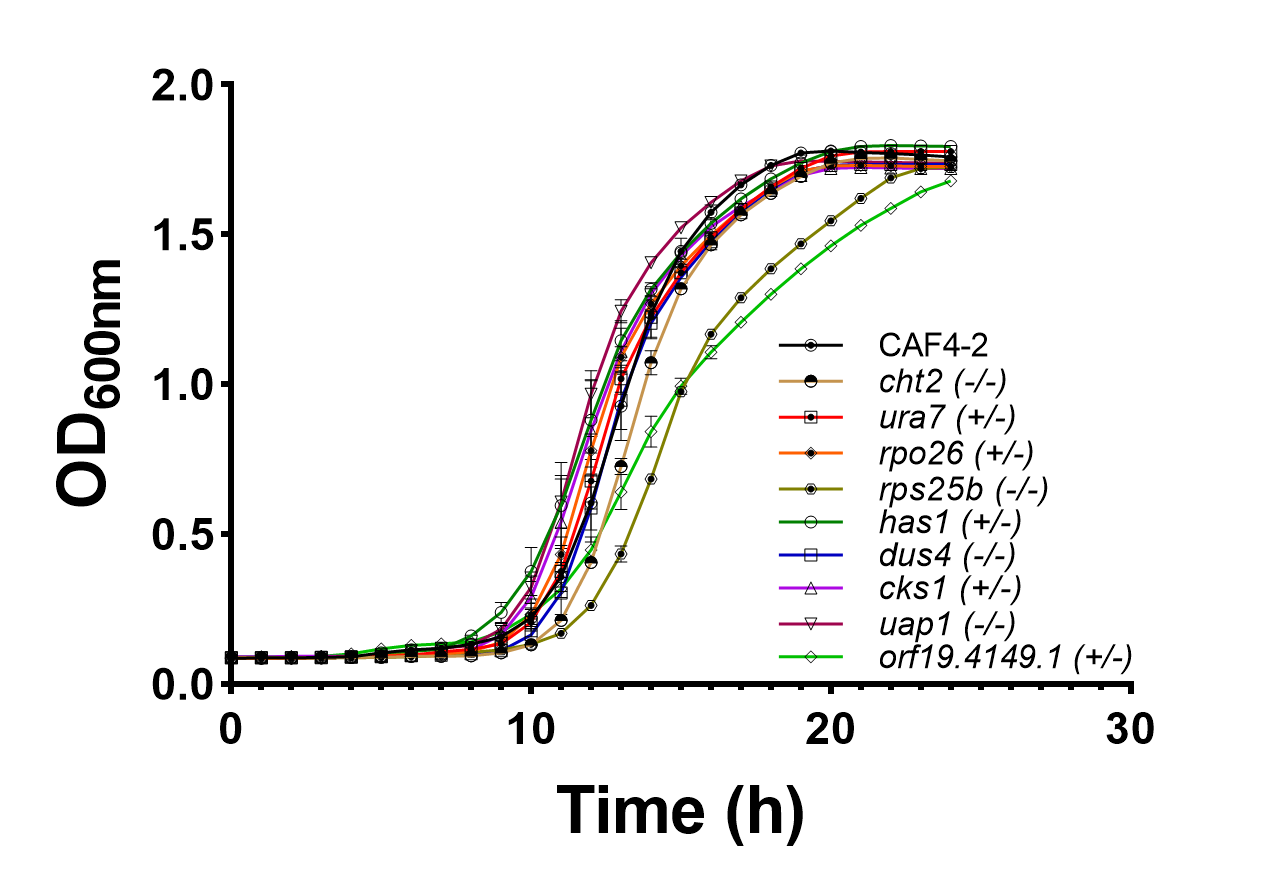

**FIG S1.** Growth curves of *C. albicans* deletion mutants vs parental CAF4-2. The panel shows parental strain CAF4-2 and Ch5 independent null mutants lacking *CHT2*, *DUS4*, *RPS25B*, *UAP1* and independent mutants lacking one copy of *URA7*, *RPO26*, *HAS1*, *CKS1* or orf19.4149.1. The cell growth was conducted in YPD medium at 35°C. Optical density was measured at 600 nm and plotted against time. The experiment was performed on three biological replicates with three technical replicates. The mean and standard deviation of optical density shown in the representative data were calculated from one biological replicate.
